# Keap1-resistant ΔN-Nrf2 isoform does not translocate to the nucleus upon electrophilic stress

**DOI:** 10.1101/2022.06.10.495609

**Authors:** Sara Mikac, Alicja Dziadosz, Monikaben Padariya, Umesh Kalathiya, Robin Fahraeus, Natalia Marek-Trzonkowska, Elżbieta Chruściel, Zuzanna Urban-Wójciuk, Ines Papak, Łukasz Arcimowicz, Tomasz Marjanski, Witold Rzyman, Alicja Sznarkowska

**Affiliations:** International Centre for Cancer Vaccine Science, University of Gdansk, ul. Kładki 24, 80-822 Gdańsk, Poland; Inserm UMRS1131, Institut de Génétique Moléculaire, Université Paris 7, Hôpital St. Louis, F-75010 Paris, France; RECAMO, Masaryk Memorial Cancer Institute, Zluty kopec 7, 65653 Brno, Czech Republic; Department of Medical Biosciences, Building 6M, Umeå University, 901 85 Umeå, Sweden; Department of Family Medicine, Medical University of Gdańsk, Gdańsk, Poland; Department of Medical Immunology, Medical University of Gdańsk, Gdańsk, Poland; Department of Thoracic Surgery, Medical University of Gdansk, Gdansk, Poland

**Keywords:** Nrf2, Nrf2 isoform 2, Keap1, NSCLC

## Abstract

The Nrf2 pathway is an essential defense pathway in a cell. It responds to oxidative and electrophilic stress via derepression of Nrf2 from Keap1-Cul3-mediated degradation, accumulation of Nrf2 in the nucleus and transcriptional activation of a number of detoxifying and cell protective Nrf2 target genes. Here we report that normal and cancer cells also express the N-terminally truncated Nrf2 isoform (ΔN-Nrf2), which originates from an alternative promoter. Co-immunoprecipitation together with molecular dynamics simulation showed that the binding between ΔN-Nrf2 and Keap1 is impaired, resulting in the much higher stability of this form. ΔN-Nrf2 is retained in the cytoplasm in response to electrophilic stress, indicating that it does not regulate transcription under the same stress stimuli as the full-length Nrf2. Altogether this data suggests that Nrf2 has other functions in cells than transcriptional activation of genes, which most probably rely on the protein-protein interactions in the cytoplasm. The regulation between these functions takes place on the level of transcription.

**Significance Statement:** This work signifies the importance of alternative transcription in assigning the function to the produced protein. Nrf2 transcripts produced from the second promoter of the Nrf2 gene give rise to the N-terminally truncated Nrf2 form (ΔN-Nrf2), which is retained in the cytoplasm upon stress, thus it has a different role in cells than transcriptional regulation. ΔN-Nrf2 is resistant to the Keap1-Cul3 degradation pathway and is highly expressed in all tested cell types. This work points to the new, cytoplasmic role of Nrf2 in cells, determined at the level of transcription.

## Introduction

Nrf2 is a stress-induced transcription factor, considered the main defense factor in the cell and a major regulator of cell survival (1). Nrf2 levels are kept low under homeostatic conditions mainly via the Keap1-Cul3 E3 ubiquitin ligase complex which mediates constitutive degradation of the Keap1-bound Nrf2 (2). Oxidative or electrophilic stress activates Nrf2 via de-repression - it modifies distinct cysteine residues in Keap1 resulting in conformational change in the complex that impairs Nrf2 ubiquitination (3, 4). In consequence, freshly synthesized Nrf2 can accumulate in the nucleus and induce expression of a battery of cytoprotective genes that ameliorate stress. Many studies revealed various functions of Nrf2 that go beyond its redox-regulating capacities, including regulation of metabolism, inflammation, autophagy, proteostasis and unfolded protein response, particularly in the context of carcinogenesis (1, 5–9).

The cytoprotective responses elicited by Nrf2 are desired in tumor cells and an aberrant Nrf2 activation has been observed in various cancer types, including the non-small cell lung cancer (10). Unregulated Nrf2 can drive carcinogenesis by activating detoxifying and antioxidative enzymes and through metabolic reprogramming (11). Constitutive Nrf2 activation in tumors is achieved via several mechanisms. One of them are point mutations in Nrf2 or Keap1 genes, which impair their binding and stabilize Nrf2 in cells (12). Another mechanism is alternative splicing. The Nrf2 gene (*NFE2L2*) produces several different transcripts that differ in promoter usage and splicing sites. Transcript 1 is the longest Nrf2 transcript that consists of five exons and gives rise to the full-length Nrf2 isoform 1, which undergoes Keap1-mediated degradation under no stress. Alternative splicing of this transcript, produces isoforms that miss exon 2, exon 3 or both. They were identified in the 6% of TCGA tumors carrying wild type Nrf2 (13). Since the Keap1 binding site is localized within exon 2, produced Nrf2 isoforms do not bind Keap1 and are resistant to the Keap1-Cul3 degradation pathway. They accumulate in the nucleus and activate expression of Nrf2 targets.

Interestingly, *NFE2L2* can be expressed from a second promoter (P2), localized downstream. P2 transcripts miss part of the open reading frame (ORF) from exon 1, encoding 16 amino acids. Recently the long-read RNA sequencing (Iso-seq) identified the Nrf2 transcript originating from P2 as the second highest expressed Nrf2 transcript under homeostatic and stress-induced conditions but the presence of encoded protein has not been reported so far (14).

Here we show that the protein product of the P2 transcripts, namely ΔN-Nrf2, is expressed in all tested lung cells including NSCLC cell lines, primary cells and normal lung fibroblasts. The lack of the first 16 amino acids impairs its binding with Keap1 and stabilizes it in cells. Interestingly, ΔN-Nrf2 does not translocate to the nucleus in response to electrophilic stress, meaning it has a different function in cells than transcriptional activation.

## Results

We have used three NSCLC cell lines: an adenocarcinoma A549, a squamous cell carcinoma RERF-LC-AI (further referred as RERF) and H1299, that differ in the Nrf2 level and activation status. A549 cells have a homozygous *KEAP1* mutation (G333C) that disrupts its binding to Nrf2, leading to the accumulation and constitutive activation of Nrf2 (15). Moreover, A549 cells have the trisomy of the chromosome 2 with *NRF2* gene, additionally causing high Nrf2 level (16). The RERF and H1299 cells do not have any known *KEAP1/NRF2* mutations (17, 18). We have also established two primary NSCLC cell lines from lung tumors of two patients (further referred as NSCLC 1 and NSCLC 2). Normal fibroblasts from the lung (further referred to as NLF) have been used as control cells.

### A stable form of Nrf2 is expressed in lung cells under homeostatic conditions

Endogenous Nrf2 migrates in 8% SDS-PAGE as three bands in all NSCLC cell lines, primary cells and normal lung fibroblasts (**Fig. 1A, *Fig. S1A***) and this pattern is detected by two different anti-Nrf2 antibodies (***Fig. S1B***). Nrf2 gene knockdown with a pool of anti-Nrf2 siRNAs reduced the expression of these forms, indicating they all represent Nrf2 species (**Fig. 1B**). Interestingly, the fastest migrating band (Nrf2*,) of approx. ∼105 kDa showed an unexpected stability upon translation inhibition compared with two other Nrf2 bands **(Fig. 1C**). It was still detected after 2 h chase with emetine in H1299 cells and after 4 h chase in NSCLC cells (**Fig. 1C** and ***Fig. S2***). Such a high stability under no stress indicates that this form might not be constitutively degraded *via* the Keap1-Cul3 system. The middle Nrf2 form had the lowest stability and in H1299 and NSCLC2 cells was not detected already after 15 min of translation inhibition. The top Nrf2 form (∼130 kDa) was still present at that time indicating higher stability than the middle form.

**Figure 1.**
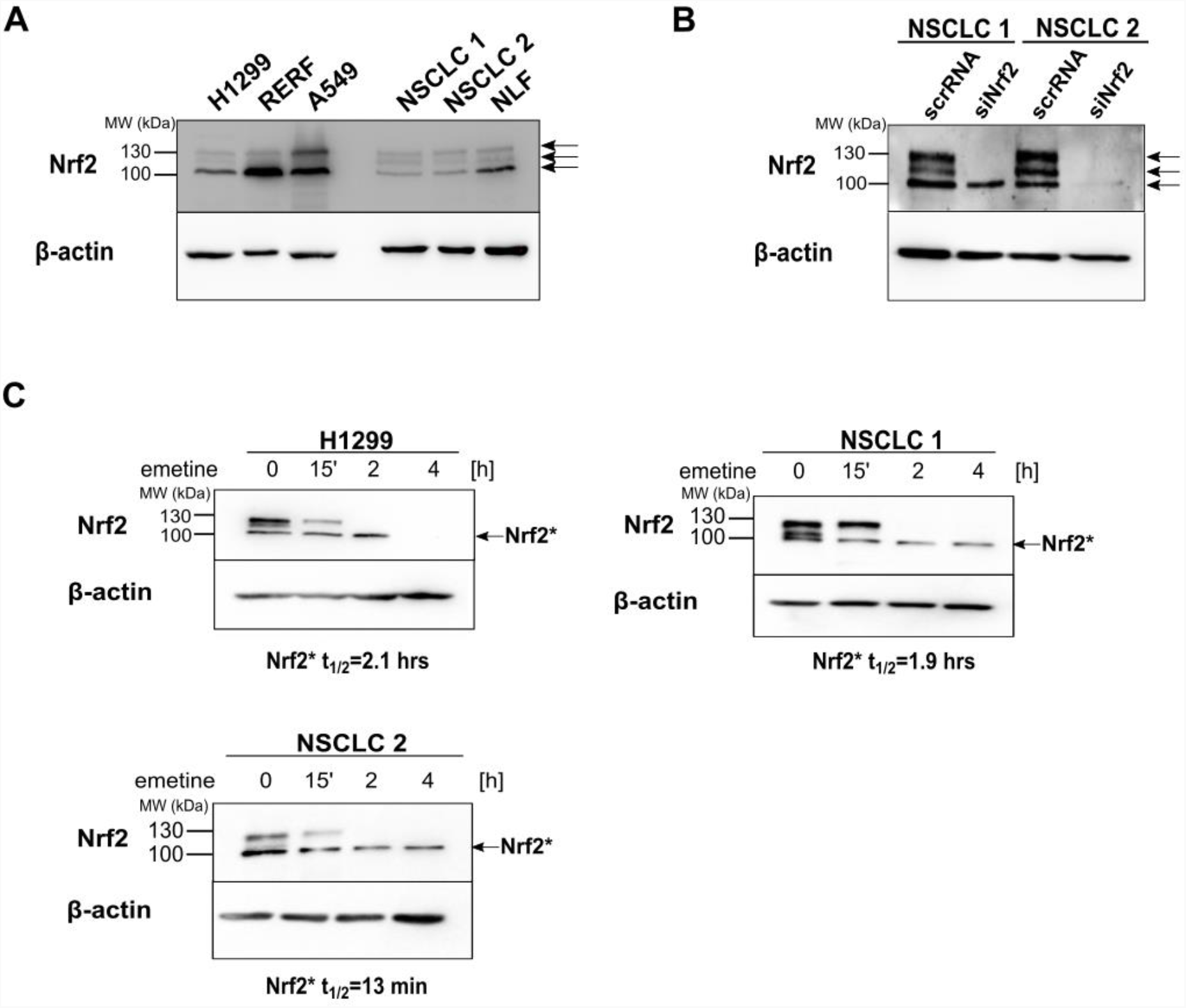
Stable Nrf2 form is expressed in lung cells under homeostatic conditions. (A) Western blot analysis of Nrf2 migration in 8% SDS-PAGE in H1299, RERF, A549 cell lines, NSCLC primary cells (NSCLC 1 and NSCLC 2) and normal lung fibroblasts (NLF). Arrows indicate three Nrf2 forms detected. (B) Nrf2 knockdown with a pool of Nrf2-targeting siRNAs (siNrf2, 25 nM) and unspecific control RNAs (scrRNA) for 48 hours in NSCLC primary cells. (C) Western blot analysis after treatment with translation elongation inhibitor emetine (20 μM) at indicated time points. Half-life of a stable Nrf2 form (Nrf2*) was calculated using nonlinear regression and a one-phase exponential decay equation based on the average band densities for each time point that were normalized to the time point zero (SI Appendix Fig. S2 and Tables S1-S3).

### The stable Nrf2 form represents N-terminally truncated Nrf2 isoform 2

To decipher the identity of detected Nrf2 forms, we first addressed if they resulted from the post-translational modifications of Nrf2. Phosphorylation is the predominant Nrf2 modification increasing its stability (19) and various Nrf2 residues are phosphorylated (20, 21). Thus we made use of lambda protein phosphatase (λPP) which removes phosphate groups from phosphorylated Ser, Thr and Tyr residues and observed changes in Nrf2 migration upon λPP (**Fig. 2A**). Upon dephosphorylation, the slowest migrating Nrf2 band disappeared while the middle Nrf2 form accumulated indicating that the top Nrf2 form is a phosphorylated Nrf2 form, that becomes reduced to the middle, unphosphorylated Nrf2 form upon λPP treatment. That is consistent with the observed stability of these forms - phosphorylated Nrf2 is more stable than the non-phosphorylated Nrf2 (**Fig. 1C**). The molecular weight (MW) of the stable Nrf2* form was not changed upon λPP treatment meaning it is not phosphorylated under homeostatic conditions. Similar results of the Nrf2 dephosphorylation were obtained with different anti-Nrf2 antibodies, recognising another Nrf2 epitope, indicating that the observed migratory pattern of Nrf2 species is not an antibody artifact (***Fig. S3***).

**Figure 2.**
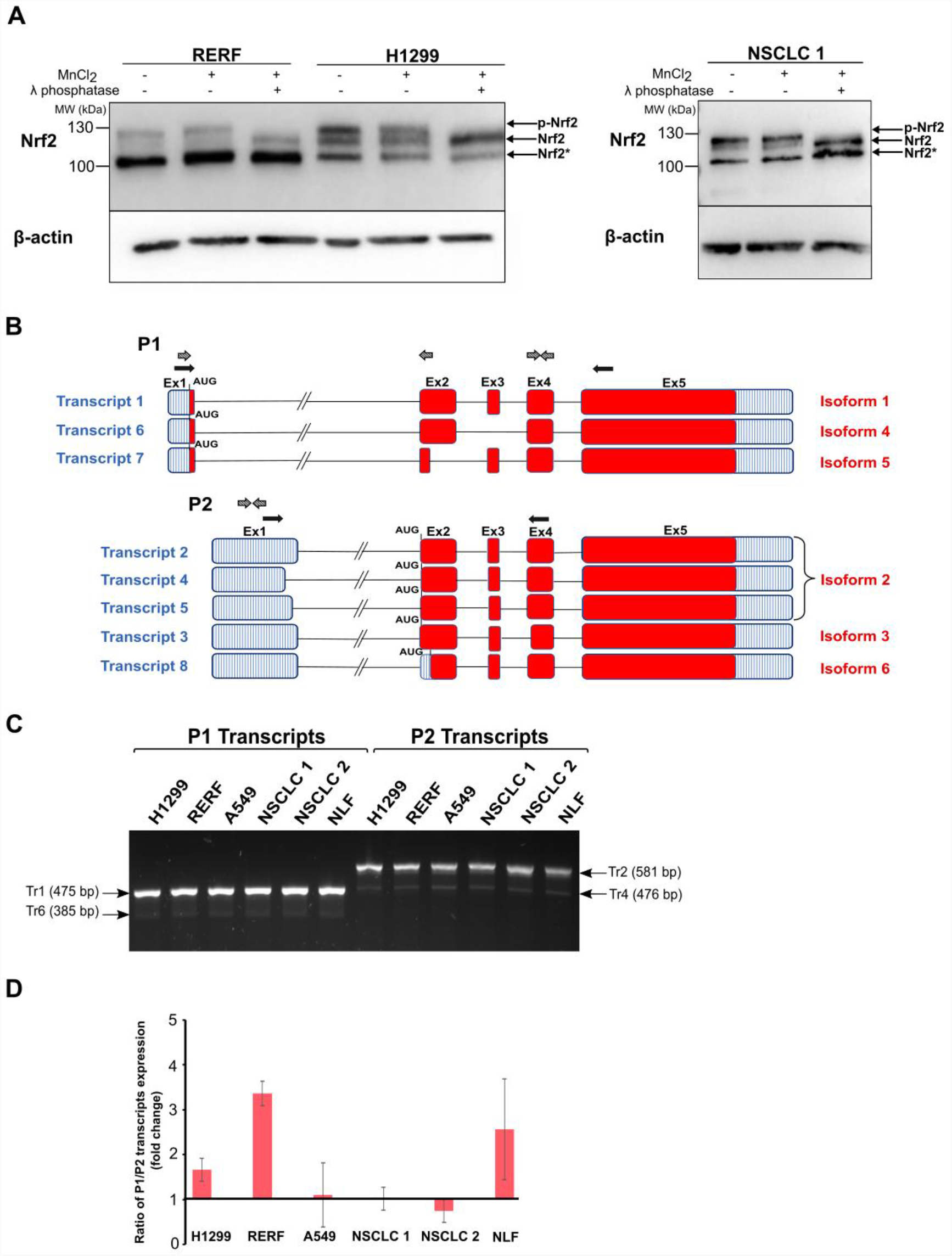
Stable Nrf2 form is not phosphorylated and originates from P2 transcript variants. (A) Lambda protein phosphatase (λPP) treatment of RERF and H1299 lysates. Cell lysates were incubated with or without λ phosphatase for 30 min in 30°C in the presence of MnCl_2_ and Nrf2 was detected by western blot. Arrows indicate different Nrf2 forms: ← p-Nrf2 for phosphorylated Nrf2; ← Nrf2 for dephosphorylated Nrf2, ←Nrf2* for a stable Nrf2 form. (B) A scheme of all possible transcript variants of Nrf2 gene (*NFE2L2*) and corresponding proteins according to NCBI. Transcripts are shown in vertical blue lines and protein encoding regions are in red. Black arrows represent primers used in reverse transcriptase PCR (RT-PCR) to identify which Nrf2 transcripts are expressed in tested cells. Dashed arrows show localization of primers for RT-qPCR for analysis of expression level of identified transcripts. P1-promoter 1; P2-promoter 2; Ex-exon. (C) Agarose gel electrophoresis (2%) after RT-PCR shows PCR products corresponding to Nrf2 transcripts (produced from P1 and P2) expressed in tested lung cells. Arrows indicate identified transcripts based on the product length (Table S4). (D) Ratio of expression of P1 vs. P2 transcripts in tested lung cells analyzed with RT-qPCR (mean ± SD, n = 3 independent experiments).

We have assumed that the stable Nrf2 form might represent a product of an alternative transcription or/and translation, especially that Nrf2 isoforms resistant to Keap1-Cul3-mediated degradation have been described in cancers before (13). According to the NCBI database, *NFE2L2* gene is expressed from two promoters which produce eight Nrf2 transcripts. Promoter 1 (P1) gives rise to transcript one, six and seven and promoter two (P2) produces transcript two, three, four, five and eight (**Fig. 2B**). Transcripts coming from P2 utilize different AUG to initiate translation than transcripts coming from P1, localized 5’ downstream to the first AUG, thus protein products of P2 transcripts are N-terminally truncated. To the best of our knowledge, only transcripts coming from P1 promoter were shown to be translated (13), while Nrf2 protein produced from P2 transcripts have not been reported so far. To study which Nrf2 transcripts are expressed in lung cells used in this study, we have designed two sets of primers, one for the P1 and second for the P2 transcripts, which amplify products of all eight Nrf2 transcripts in the reverse transcriptase PCR (RT-PCR) (**Fig. 2B** and Table S4). We separated these products in 2% agarose to differentiate them by size and found that the most abundant transcripts expressed in all cell types are transcript 1 and transcript 2 (**Fig. 2C**). Transcript 1 represents the full-length Nrf2 transcript and is translated to the full-length Nrf2 isoform 1 of 605 amino acids. Transcript 2 is transcribed from the P2 promoter and is translated to the isoform 2 which lacks the first 16 amino acids due to the utilization of the downstream AUG. This analysis also revealed the presence of transcript 6 expressed from the P1 and transcript 4 from the P2, though produced on a much lower scale (**Fig. 2B**). Transcript 6 undergoes alternative splicing, in which exon 3 is spliced out. Transcript 4 differs from transcript 2 in the 5’UTR sequence and gives rise to the same protein, Nrf2 isoform 2 (**Fig. 2B**). The end point RT-PCR gives us qualitative information about which transcripts are expressed but cannot provide precise quantitative information on their expression levels. Thus we made use of quantitative RT-PCR (qPCR, Real-time PCR) and analysed the ratio of expression of P1 to P2 transcripts in all tested lung cells (primers used for qPCR are presented in Table S5). We found that the majority of cells expressed the same amounts of P1 and P2 transcripts, only RERF and fibroblasts produced slightly more P1 transcripts (**Fig. 2D**). From this analysis we could conclude that Nrf2 isoform 2, produced from transcripts 2 and 4, could represent the fastest migrating, 105 kDa Nrf2 form, detected in Western blot. We have named it ΔN-Nrf2 due to the N-terminal truncation. The 16 aa-longer, full-length isoform 1 migrated just above the isoform 2, as the middle band. The slowest migrating Nrf2 form was the phosphorylated isoform 1, which after dephosphorylation was reduced to the non-phosphorylated isoform 1 (middle band, **Fig. 2A**).

### Binding of ΔN-Nrf2 isoform to Keap1 is impaired

Next question was why the lack of first 16 amino acids in ΔN-Nrf2 isoform would have such a profound impact on the Nrf2 stability. Since the full-length Nrf2 (isoform 1) is rapidly degraded under no-stress conditions via Keap1-Cul3 ubiquitin ligase, the reason for an increased stability of the ΔN-Nrf2 should be the resistance to this degradation pathway. We therefore assumed that binding of ΔN-Nrf2 to Keap1 was impaired. To validate this hypothesis, we made use of molecular modeling and molecular dynamics simulation (MDS) that helped us understand the structural differences between isoforms and their impact on the structural dynamics of the Keap1 binding. We first built a model of the full-length Nrf2-Keap1 complex based on the available crystal structure of their binding domains (**Fig. 3A, *Fig. S3A***). This model showed that the first 16 amino acids of Nrf2 constituted a part of the first alpha-helix of Nrf2. Next, we performed MD simulation to investigate the binding affinity and structural dynamics of the Nrf2-Keap1 complex in the course of time. MDS revealed that the first 16 aa of Nrf2 built a protein ‘tail’ which *via* its constant movements was stabilizing a closed conformation of Nrf2 and sustaining interaction with Keap1. When these amino acids are missing, Nrf2 undergoes conformational change and opens up, which destabilizes interaction with Keap1 (**Fig. 3B, *Fig. 3SA***,***B***), resulting in a decreased number of hydrogen bonds between ΔN-Nrf2 isoform and Keap1 (**Fig. 3C**). We further aimed to verify differences in Keap1 binding between Nrf2 isoforms in cells. We have chosen RERF cells as under no-stress they express high levels of ΔN-Nrf2 isoform and a very low amount of full-length Nrf2. We have immunoprecipitated total Nrf2 and analyzed the amount of Nrf2-bound Keap1 under homeostatic conditions and after a specific small molecule inhibitor of neddylation, MLN4924, which blocks E3 ubiquitin ligase activity of cullins and leads to the accumulation of their substrates, such as Nrf2 (22, 23) (**Fig. 3D**). Interestingly, inhibition of Cul3 activity with MLN4924 led to the accumulation of full-length Nrf2, but not ΔN-Nrf2 isoform, indicating that ΔN-Nrf2 is not regulated via the cullin system (**Fig. 3E, *Fig. S3D***). The amount of Keap1 bound to Nrf2 was very little under no-stress, consistent with a constitutive degradation of full-length Nrf2, but after MLN4924 treatment it increased dramatically (**Fig. 3E, *Fig. S3E***). Since the amount of ΔN-Nrf2 has not changed, this increase is due to the Keap1 binding to the full-length Nrf2. Consistently, knockdown of Keap1 in primary NSCLC cells resulted in the accumulation of phosphorylated Nrf2, but did not affect the amount of ΔN-Nrf2 isoform (**Fig. 3F**).

**Figure 3.**
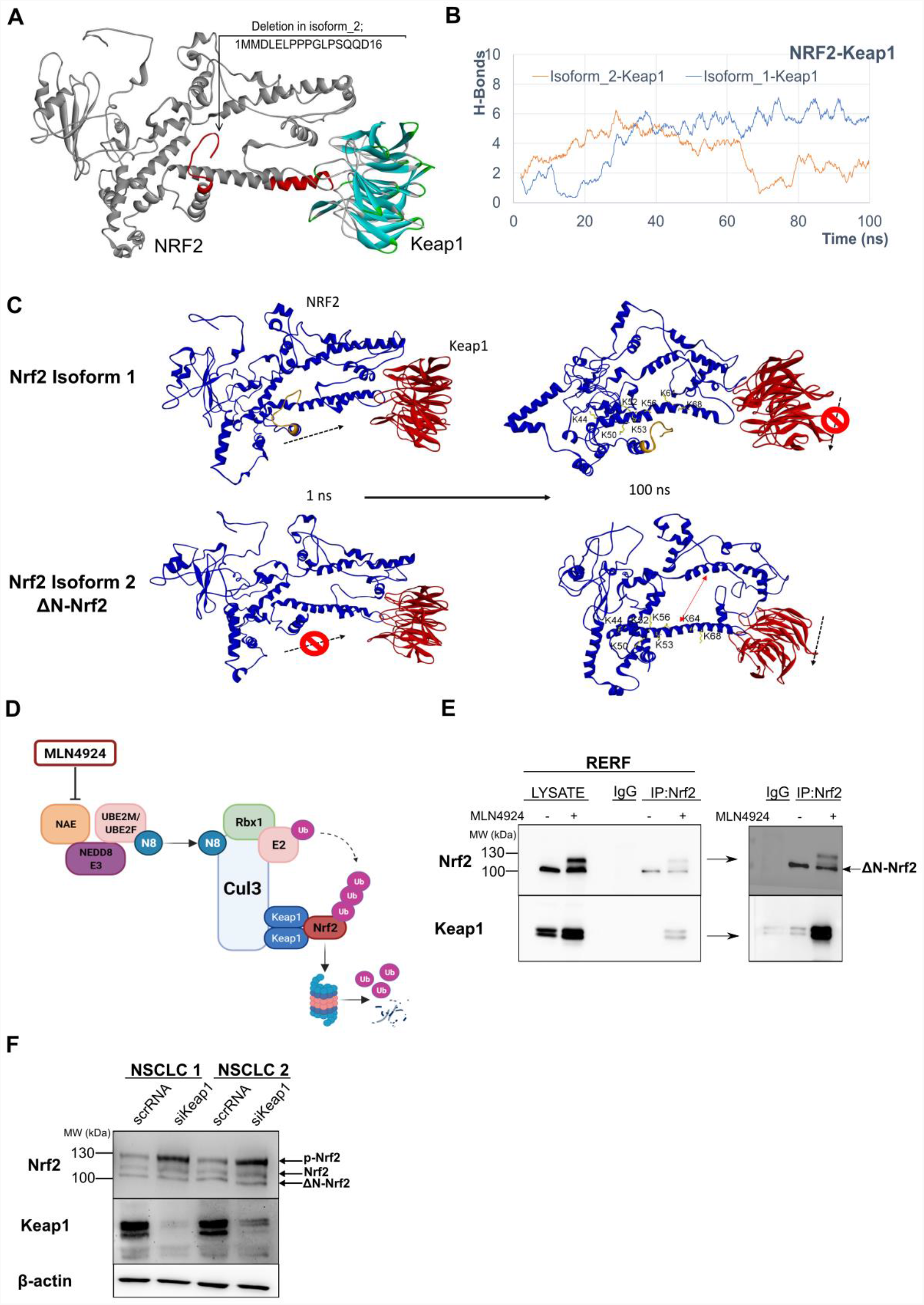
Binding of ΔN-Nrf2 to Keap1 is impaired. (A) A created model of the Nrf2 protein bound to Keap1. An arrow indicates the first 16 aa missing in the Nrf2 isoform_2 (ΔN-Nrf2). The Nrf2 high affinity binding motif (ETGE) interacting with Keap1 is marked in red (B) Results of molecular dynamics simulations (MDS) showing a number of hydrogen bonds created during 100 ns MD simulation between Keap1 and Nrf2 isoform_1 (blue line) and Keap1 and Nrf2 isoform_2 (red line). (C) Structures of Nrf2-Keap1 complexes before and after 100 ns MD simulations. The first 16 aa of Nrf2 isoform_1 build a ‘tail’ (in gold) which keeps the closed conformation of Nrf2 and stabilizes binding with Keap1 in time. The structure of Nrf2 isoform_2 opens up during MDS which destabilizes Keap1 binding. The lysine residues of Nrf2 that undergo Keap1-Cul3-mediated ubiquitination are indicated (D) The mode of action of MLN4924 neddylation inhibitor on the Cul3-Keap1-E3 ligase complex. The Cul3-Keap1-E3 ligase complex is active in a neddylated state, resulting in Nrf2 ubiquitination and degradation. The complex is inactivated by a neddylation inhibitor MLN4924, which binds to NAE, blocks its enzymatic activity and inhibits neddylation. It results in the accumulation of ubiquitinated Nrf2 in the cytoplasm. Abbreviations: CUL3 cullin 3; E2, ubiquitin-conjugating enzyme; N8, NEDD8; NAE, NEDD8-activating enzyme; NEDD8, NEDD8 ubiquitin-like modifier; RBX1, RING-box 1; UBE2M, ubiquitin-conjugating enzyme E2M; UBE2F, ubiquitin-conjugating enzyme E2F; Ub, ubiquitin. Created with Biorender.com. (E) Western blot analysis of RERF cells after Nrf2 precipitation (IP) under steady-state and upon treatment with a neddylation inhibitor MLN4924 for 12 hours. Precipitates were probed with anti-Keap1 antibodies to analyze levels of Nrf2-bound Keap1. MLN4924 induced the full-length Nrf2 and p-Nrf2, but not the ΔN-Nrf2 levels. Under steady-state, little Keap1 was bound to Nrf2 but upon MLN4924 the amount of Nrf2-bound Keap1 was largely increased. The right side panel presents the co-IP results after longer exposure. Arrow indicates Nrf2 isoform 2 (ΔN-Nrf2). (F) Keap1 knockdown with Keap1-specific siRNAs (Santa Cruz Biotechnology, 25 nM) in NSCLC primary cells induced phosphorylated Nrf2 and non-phosphorylated Nrf2, but not the ΔN-Nrf2 form. Arrows indicate different Nrf2 forms: ←p-Nrf2 for phosphorylated Nrf2; ←Nrf2 for non-phosphorylated Nrf2; ←ΔN-Nrf2 for the N-terminally truncated Nrf2 isoform 2.

### ΔN-Nrf2 isoform does not translocate to the nucleus upon electrophilic stress

Finally we wanted to learn what is the role of the ΔN-Nrf2 and how it responds to stress. We have used an electrophilic food preservative tert-butylhydroquinone (tBHQ), known to stabilize Nrf2 and to induce its nuclear transfer (24–26). tBHQ activates Nrf2 both in a Keap1-dependent fashion via triggering Keap1 degradation (27), and in the Keap1-independent way by increasing intracellular free zinc concentration which inhibits phosphatases activity (28) and induces Nrf2 phosphorylation (20). tBHQ treatment induced both unphosphorylated and phosphorylated full-length Nrf2, while ΔN-Nrf2 was not induced in any of the cells analyzed (**Fig. 4A**). We have found that ΔN-Nrf2 was localized in the cytoplasm both under homeostatic conditions (**Fig. 4B**) and after tBHQ (**Fig. 4C**) in cell lines, primary NSCLC cells and normal fibroblasts, while phosphorylated Nrf2 isoform 1 was detected mainly in the nucleus under both normal conditions and, more abundant, after tBHQ in all cell types (**Fig. 4B** and **C**). These results suggest that a stable ΔN-Nrf2 isoform does not have a transcription factor function but rather plays a role in cytoplasm where it can be engaged in the protein-protein interactions.

**Figure 4.**
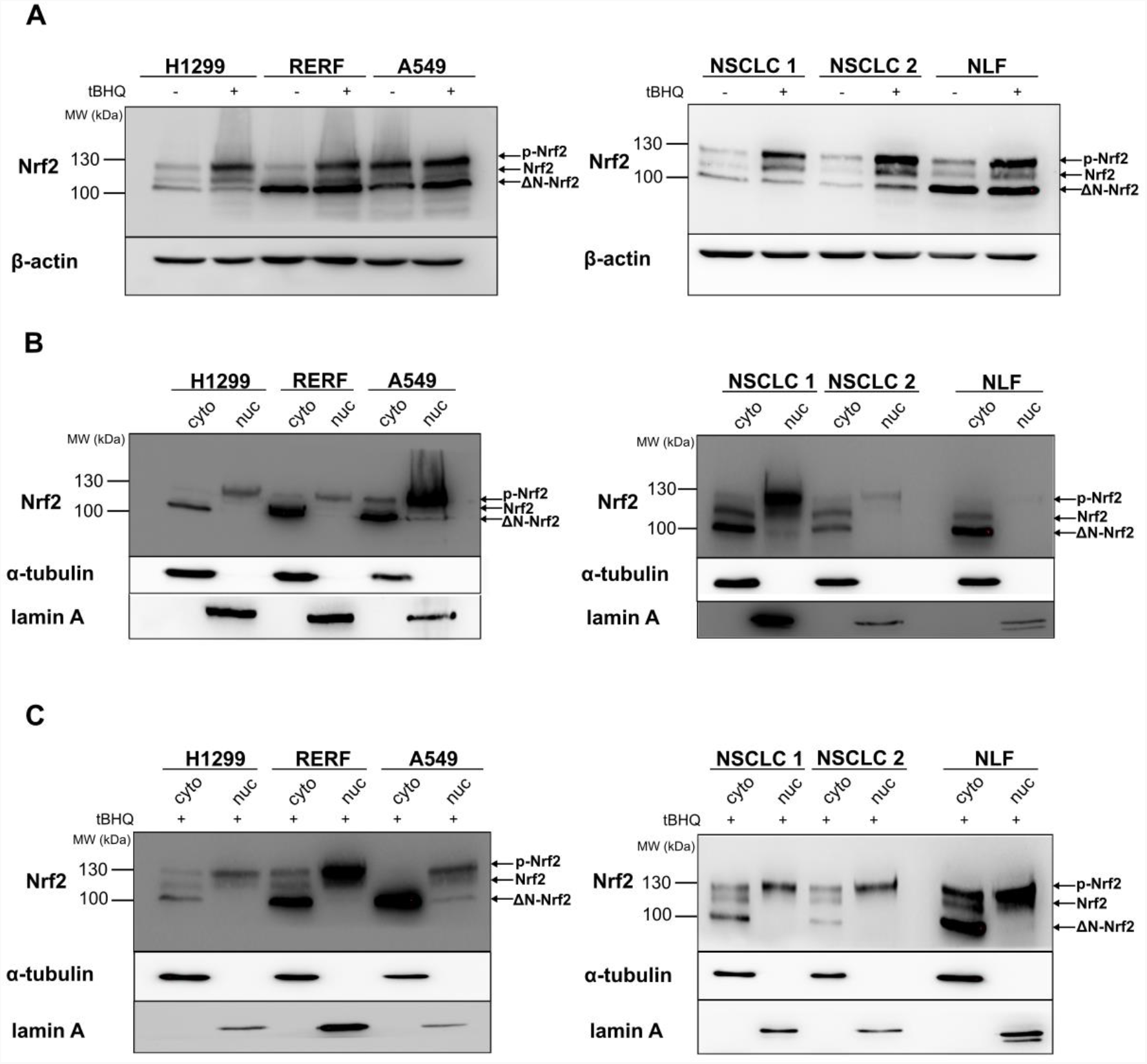
ΔN-Nrf2 isoform does not translocate to the nucleus in response to electrophilic stress. Left panels present results in cell lines: H1299, RERF and A549, right panels present results in primary NSCLC cells (NSCLC1, NSCLC2) and normal lung fibroblasts (NLF). (A) Western blot analysis of cell lysates after tBHQ treatment (20 uM) for 6 hours. (B) Cellular fractionation under steady state conditions. Lamin A was used as a nuclear marker, α-tubulin was used as a cytoplasmic marker. (C) Cellular fractionation after tBHQ treatment (20 uM) for 6 hours. Lamin A was used as a nuclear marker, while α-tubulin was used as a cytoplasmic marker. Arrows indicate different Nrf2 forms: ←p-Nrf2 for phosphorylated Nrf2; ←Nrf2 for non-phosphorylated Nrf2; ←ΔN-Nrf2 for the N-terminally truncated Nrf2 isoform 2.

## Discussion

The Nrf2 pathway is associated with a stress-induced activation of a cell-protective gene expression. This study indicates that the Nrf2 role might go beyond gene regulation. The here-identified N-terminally truncated Nrf2 isoform 2 (named ΔN-Nrf2) is not degraded by the Keap1-Cul3 complex under no stress and it does not translocate to the nucleus to regulate gene expression. Electrophilic stimulation with t-BHQ did not induce its nuclear transfer either. ΔN-Nrf2 originates from the second promoter of *NFE2L2* gene, localized downstream to the first one, and lacks the first 48 base pairs of the ORF sequence compared to the full-length Nrf2 (isoform 1). Molecular dynamics simulation and co-immunoprecipitation studies show that it exhibits an impaired binding with Keap1 and in consequence is more stable and more abundant than isoform 1 under no stress. Inhibition of cullins activity by a specific small molecule neddylation inhibitor MLN4924 did not result in the accumulation of ΔN-Nrf2, meaning that this isoform is not degraded via the cullin pathway. This is the first indication of a degradation-resistant Nrf2 form whose role in cells is other than the transcription factor.

There are many types of cancers that show intrinsically high Nrf2 activity, referred to as Nrf2-addicted cancers (29). The Nrf2 pathway is dysregulated there via point mutations or alternative splicing of *NFE2L2* gene, which impair the binding with Keap1. Produced Nrf2 species are highly stable and actively regulate expression of Nrf2 targets. Thus, an escape from degradation was considered sufficient to maintain a constitutive expression of Nrf2 targets by Nrf2 mutants or spliced variants. But ΔN-Nrf2 did not translocate to the nucleus, both under homeostatic and upon electrophilic stress conditions with an exception of A549 cells where a fraction of this form was observed also in the nucleus (Fig. 4B and C). A549 is an example of the Nrf2 addicted cancer celI line with unregulated Nrf2 levels mainly due to Keap1 mutations. Perhaps this dysregulation affects the expression of all the *NFE2L2* transcripts via Nrf2 autoregulatory feedback loop, resulting in some ‘leakage’ of ΔN-Nrf2 to the nucleus. So far there is no data on the putative nuclear localization signal (NLS) that would be contained within the first 16 amino acids of Nrf2, thus missing in the ΔN-Nrf2 isoform. The closest functional NLS has been mapped to the residues 42-53 (30) and the two others reside within C-terminus of the protein (515–518 and 587–593 amino acids) (30, 31). It means that ΔN-Nrf2 retains all the NLS present in the full-length Nrf2. Another possible explanation of the cytoplasmic localization of ΔN-Nrf2 is that Nrf2 needs to be post-translationally modified to enter the nucleus and that such a modification is missing in the ΔN-Nrf2. It was suggested before that Nrf2 has to be phosphorylated to translocate to the nucleus (32). Phosphorylation of Ser 40 was the best studied and despite it promoted dissociation from Keap1, it was not required for the accumulation in the nucleus and transcriptional activation of Nrf2 targets (33, 34). In line with this, we have observed that nuclear fractions contained both the non-phosphorylated and the phosphorylated full-length Nrf2 while the non-phosphorylated ΔN-Nrf2 was detected almost exclusively in the cytoplasm both under homeostatic conditions and upon electrophilic stress (Fig. 4C). We cannot exclude the presence of yet unknown modification within the first 16 amino acids of Nrf2 that regulates its nuclear shuttling.

Recently Otsuki et al. studied transcript variants of Keap1-Nrf2 pathway utilized in the oxidative stress response with the use of the long read RNA isoform sequencing (Iso-Seq) (14). Authors did identify the ΔN-Nrf2-encoding transcript as the second highest expressed, after the isoform 1 transcript, under both normal and stress-induced conditions and their expression ratio did not change upon stress-induction. Despite this, the presence of the encoded protein has not been reported so far. This work for the first time reports the detection of the Nrf2 isoform 2 (ΔN-Nrf2) in various cancer and normal cells. We have tested five different cell lines and in each of them the ΔN-Nrf2 isoform was expressed similarly to the full-length Nrf2 on the transcript level and was even more abundant than isoform 1 on the protein level. This difference most probably results from the different stability of the two isoforms but variations in the translation rate are also possible.

It seems peculiar why the existence of the stable ΔN-Nrf2 isoform has not been detected before, since from what we have observed it is expressed in every lung cell line tested here as well as in cells from different tissues (unpublished observation). The answer might lie in the challenges in Nrf2 detection in Western blot (WB): (1) it does not migrate according to the predicted molecular weight; (2) it migrates as a few bands which was attributed to post-translational modifications; (3) for a long time anti-Nrf2 antibodies have not been very specific; (4) in stability studies, Nrf2 was ectopically expressed from vectors carrying the full-length isoform 1 cDNA). Nowadays, thanks to the specific and sensitive antibodies available on the market and with the proper protein separation in the 8% SDS-PAGE, the identification and characterisation of endogenous Nrf2 isoforms is possible. We have reported the existence of a stable Nrf2 form before (35), but these studies allowed us to recognise its alternative transcription origin.

The big question to be addressed by future studies is what is the function of ΔN-Nrf2 in the cytoplasm and if it associates with carcinogenesis. So far the Nrf2 pathway has been studied solely from the perspective of Nrf2-regulated genes, but the identification of the cytoplasmic Nrf2 form will surely redirect this research towards the protein-protein interactions and their consequences. Other essential transcription factors (TFs) such as p53, p63 and p73 also possess alternatively transcribed and translated isoforms with altered functions affecting the whole pathway (36–38). N-terminally truncated isoforms of these TFs lost the transcription regulation potential due to a deletion in transactivating domains. Since p53 family members act as tetramers and all monomers in a tetramer have to be transcriptionally active, dimerization of a truncated with a full-length form results in the dysfunctional oligomer and decreases the transactivating potential of the pathway. Expression of these truncated isoforms is associated with various types of cancers (39) and illustrates how an alternative expression of one gene imposes an extra layer of regulation on the whole pathway. It will be very interesting to understand the cytoplasmic role of Nrf2 under steady state and stress and learn how Nrf2 isoforms act together to harmonize cell defense and protection responses.

## Materials and Methods

### Cell lines

Non-small cell lung cancer cell lines A549, RERF-LC-AI and H1299, as well as normal lung fibroblasts (NLF), were purchased from RIKEN BRC Cell Bank (Tsukuba, Ibaraki, Japan). Cancer cell lines were cultured in Dulbecco’s modified Eagle’s medium (Gibco, Thermo Fisher Scientific), with 8% of Fetal Bovine Serum (Gibco, Thermo Fisher Scientific) and 1% of Penicillin-Streptomycin (10 000 U/mL, Gibco, Thermo Fisher Scientific), while normal lung fibroblasts were cultured in Ham’s F-12 Nutrient Mix (Gibco, Thermo Fisher Scientific), with 10% of Fetal Bovine Serum (Gibco, Thermo Fisher Scientific) and 1% of Penicillin-Streptomycin (10 000 U/mL, Gibco, Thermo Fisher Scientific). Cells were maintained at 37°C under humidified conditions with 5% CO_2_.

### Establishment of the primary non-small cell lung cancer (NSCLC) cell lines

Cancer samples (0.25-1g) were obtained from NSCLC patients admitted to the Clinic of Thoracic Surgery of University Clinical Centre of Medical University of Gdańsk (MUG) for surgical resection of the primary tumour. Patients were not previously treated with any anti-cancer therapy and no metastases were detected. Samples were cut into 1-2 mm fragments and washed 3 times with PBS to remove contaminating debris and erythrocytes. Subsequently digestion solution (DS) in 1:1 of DS (ml): tissue mass (g) ratio was added. The tissue was gently agitated at 37°C until single cell suspension was obtained. DS contained 5% collagenase type I (Sigma-Aldrich; 5mg/ml), 19% of PBS and 1% of fetal bovine serum (FBS). The collagenase was inactivated with an equal volume of 10% FBS supplemented LG-DMEM medium (PAA), followed by filtration of the resulting cell suspension through a 100 μm nylon cell strainer (Falcon). Subsequently, the cells were washed with PBS. Then, erythrocytes lysis buffer was used to remove erythrocytes (10 min. incubation at room temperature, RT). After subsequent centrifugation (600g, 10 min.) the cells were washed with 4% FBS supplemented PBS and subjected for tumour infiltrating leukocyte depletion. For this purpose, positive immunomagnetic selection of CD45+ cells were performed with EasySep Human CD45 Depletion Kit II (StemCell Technologies, Canada) according to the manufacturer instructions (post-isolation purity 96-99%). Thus, untouched NSCLC and lung-derived cells were isolated. Cells were plated into 75cm2 culture flasks in TumorPlus263 medium without serum. Establishment of the primary cell line was confirmed with flow cytometry (FAP-CK19+ phenotype; FACS Aria Fusion, BD Biosciences, USA) and when continuous proliferation of the cultured cells was observed after 2-month expansion *in vitro*. Cells that fulfilled both criteria were used for the experiments.

### Lipid-mediated reverse transfection

Cells were seeded in the 6-well plates, 100,000 cells/well and transfected with control siRNA-A (ON-TARGET plusTM Control Pool, DharmaconTM, referred in text as scrRNA), as a control for transfection (25 nM), with small-interfering RNA for silencing of Nrf2 expression (siRNA, ON-TARGET plusTM SMART pool, DharmaconTM) and small-interfering RNA for silencing of Keap1 expression (siRNA, Santa Cruz Biotechnology) in concentration of 25 nM, with 4 μL/well of Lipofectamine 3000 reagent (Invitrogen, Thermo Fisher Scientific), according to manufacturer’s instructions. Western blot was performed 48 h after transfection.

### Western blot analysis

Western blot analysis was performed as described previously (35). Primary antibodies that were used are anti-NRF2 [EP1808Y] – ChIP Grade (cat. no. ab62352; Abcam), anti-NRF2 (D1Z9C) XP antibody (cat. no. 12721; Cell Signaling Technology), anti-tubulin (DM1A, Cell Signaling Technology), anti-lamin A (C-3, sc-518013; Santa Cruz Biotechnology), anti-Keap1 (cat. no. NBP2-03319; NOVUS Biologicals) and anti-β-actin (cat. no. A2228; Sigma-Aldrich).

### Cellular Fractionation

Separation of nuclei from cytoplasm was performed according to the REAP method described by Suzuki et al (40). Briefly, 4000,0000 cells were resuspended in 400 μL of ice-cold 0.1 % NP-40 in PBS by gentle pipetting and centrifuged at 500 g for 10 seconds. Supernatant (cytosolic fraction) was collected and Laemmli buffer was added to final concentration 1x. Pellets containing nuclei were washed with 300 μL of 0.1 % NP-40 in PBS, centrifuged at 500 g for 10 seconds, resuspended in 200 μL 1x Laemmli buffer and sonicated 15 min. Samples were boiled for 10 min and analyzed by western blot.

### Co-immunoprecipitation

Cells were lysed using ice-cold lysis buffer (150 mM NaCl, 25 mM Tris-HCl, 0.5% Triton x-100; pH 7.5) and centrifuged 12,000xg for 15 min. Lysates were pre-cleared with Protein G magnetic beads (Thermo Fisher) for 30 min at 4°C. Supernatants were incubated with anti-NRF2 [EP1808Y] – ChIP Grade (cat. no. ab62352; Abcam) antibody overnight at 4°C and Nrf2-antibody complexes were precipitated via 30 min incubation with beads. Beads were washed three times in PBS and proteins were eluted in 2x SDS-loading buffer, at 50°C for 10 minutes. Samples were analyzed by western blot.

### Identification of Nrf2 transcript variants via RT-PCR

Cells were seeded at 800,000 in 60 mm plates and 24 h later RNA was isolated (Qiagen RNeasy Kit). 1 ug of RNA was reverse transcribed (Applied Biosystem), diluted 2-times and 1 μl was taken for each PCR reaction (20 ul) containing: 0.02 U HotStarTaq® DNA polymerase (Qiagen, Germany), 1 x PCR buffer, 0.2 mM dNTPs mixture and 200 nM of each primer. Primers were designed in a way to enable a length-based discrimination of Nrf2 transcript variants expressed in cells (Table S4). PCR conditions used: initial heating 15 min, 95°C followed by 45 cycles of 95°C (20 s), 62°C (20 s), 72°C (60 s) and a final extension step of 72°C for 3 min. Products were separated in 2% agarose and analysed. The identity of PCR products was confirmed with Sanger sequencing.

### Evaluation of Nrf2 transcripts expression by RT-qPCR

Cells were seeded at 800,000 in 60 mm plates and 24 h later RNA was isolated (Qiagen RNeasy Kit). 1 μg of RNA was reverse transcribed (Applied Biosystems™ High-Capacity cDNA Reverse Transcription Kit), diluted 100x and 2 μl were taken for qPCR reaction. Quantification of transcript variants expression was performed according to the previously described method (41), where a product of Nrf2 gene common to all transcript variants was used as an internal reference to calculate relative expression of selected variants amplified with variants-specific primers (Table S5). Under qPCR conditions used, the reaction efficiency for each primer pair was similar (∼2). First set of primers amplified a product common for transcripts 1,6 and 7, second – a product of transcripts 2,3 and 8 and the third – product common to all Nrf2 transcript variants (Table S5).

### Molecular dynamics simulations and molecular modeling method

The structures for both isoforms (Isoform_1 and Isoform_2 with missing 1MMDLELPPPGLPSQQD16) of NRF2 were retrieved applying homology modeling modules implemented in the C-I-TASSER package developed in Zhang’s lab (42). C-I-TASSER, uses convolutional neural-network based contact-map predictions to guide the I-TASSER fragment assembly (43). To build a complete length NRF2 protein structure, the pdb id. 6gmh (44) was used as a template. These modeled NRF2 isoforms’ structures were energy minimized applying the CHARMM27 forcefield (45) implemented in the Molecular Operating Environment (MOE; Chemical Computing Group Inc., Montreal, QC, Canada) (46) package. Furthermore, the crystal structure of high affinity binding domain from NRF2 binding with Keap1 protein is available (pdb id.: 2flu (47, 48)), hence it was used as a template to define the conformation of full NRF2 protein structure with Keap1. The modeled NRF2-Keap1 complex was further energy optimized in the MOE package, applying the CHARMM27 forcefield (45). Followingly, to understand their structural properties NRF2(isoform_1)-Keap1 and NRF2(isoform_2)-Keap1 were processed by molecular dynamics simulations (MDS) approach. MDS was performed using the GROMACS 4.6.5 (49) program (GROMACS; Groningen Machine for Chemical Simulations) assigning the CHARMM27 forcefield (45, 50, 51). Each prepared model system was solvated in simple point charge (SPC) water molecules (52) and Na^+^Cl^-^ counter ions, and in a 10 Å thick dodecahedron simulation box. Periodic boundary conditions were applied and using steepest descent algorithm the systems were minimized for 50,000 steps. Particle Mesh Ewald (PME) method (53) and the LINCS algorithm (54) were used to treat electrostatic interactions (van der Waals and Coulomb interactions were set to 10 Å) and bond lengths, respectively. NPT (isobaric-isothermal) ensemble simulation was implemented to equilibrate all modeled systems, with temperature maintained at 300 K by V-rescale thermostat (54) and pressure at 1 bar and Parrinello-Rahman barostat (55). Leapfrog integrator (56) was used to perform 100 ns simulations of each system, and trajectories were saved every 10 ps. Results obtained from MDS were analyzed using GROMACS, VMD (Visual Molecular Dynamics) (57), and MOE (Chemical Computing Group Inc., Montreal, QC, Canada) / BIOVIA Discovery Studio (Dassault Systèmes, BIOVIA Corp., San Diego, CA, USA).

## Acknowledgments

The study was supported by “International Centre for Cancer Vaccine Science” that is carried out within the International Research Agendas Programme of the Foundation for Polish Science co-financed by the European Union under the European Regional Development Fund, Polish National Science Centre PRELUDIUM9 grant No. 2015/17/N/NZ3/03773 and UGrants Advanced No. 533-0B00-GA08-22.

