## Supplementary information for "Keap1-resistant ΔN-Nrf2 isoform does not translocate to the nucleus upon electrophilic stress"

**This PDF file includes:**

Supplementary text

Figures S1 to S4

Tables S1 to S5

SI References

**Supplementary Information Text**

**Materials and Methods**

*Treatment with translation inhibitor emetine dihydrochloride*

Cells were seeded in 6-well plates, 300 000 cells/well. 48 hours later cells were treated with emetine dihydrochloride (20 µM), for 15 minutes, 4, 8, 16 and 24 hours. Cells were collected and Nrf2 levels were analyzed by western blot with anti-NRF2 [EP1808Y] – ChIP Grade (cat. no. ab62352; Abcam). For calculation of the protein half-life, average band densities for each time point were normalized to controls, and data were fitted using nonlinear regression and a one-phase exponential decay equation using GraphPad Prism software.

*Treatment with neddylation inhibitor MLN4924*

Cells were seeded in 6-well plates, 500,000 cells/well. 24 hours later, cells were treated with neddylation inhibitor MLN4924 (1 μM) for 12 hours. Cells were collected and Nrf2 levels were analyzed by western blot.

*Treatment with Lambda Protein Phosphatase (λPP)*

800,000 cells were lysed in 250 µL of RIPA lysis buffer, sonicated for 15 min on ice and briefly centrifuged at 13,000 x g. For dephosphorylation, 40 µL of cell lysate was incubated with 400 U of λPP (New England Biolabs) in the dedicated buffer and in the presence of manganese ions at 30°C for 30 min. Control samples underwent the same treatment, but without the enzyme. The Nrf2 phosphorylation was analyzed by western blot.

*Treatment with tert-butylhydroquinone (tBHQ)*

Cells were seeded in 60 mm plates, 1 000 000 cells/plate. 24 hours later, cells were treated with tert-butylhydroquinone (tBHQ) (20 μM). Cells were collected after 6 hours and cellular fractionation was performed. Nrf2 levels were analyzed by western blot.

**Figures**

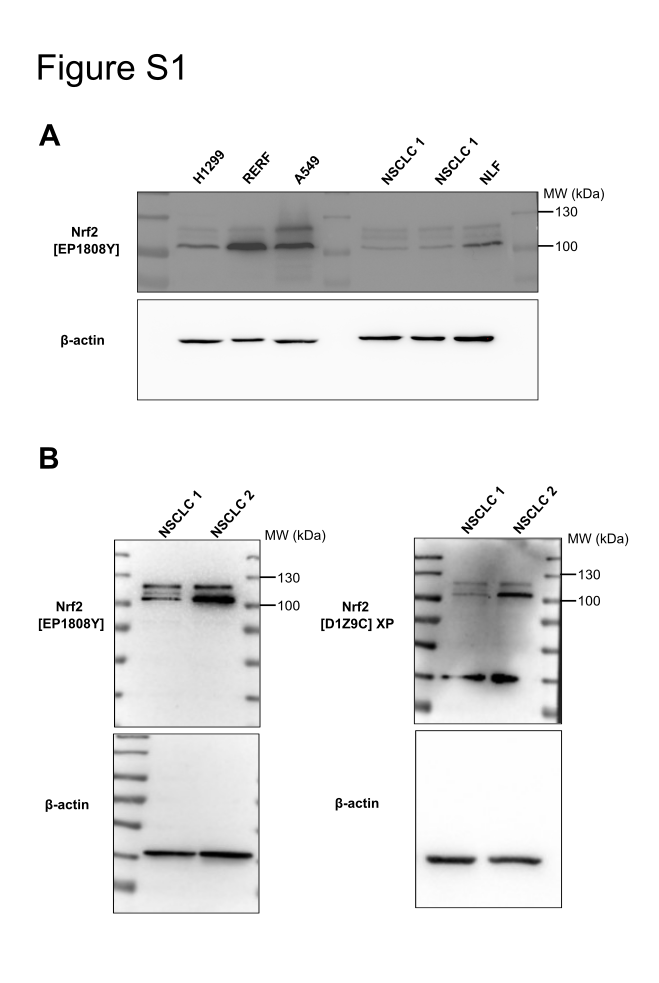

Figure S1. **Stable, non-canonically regulated 105 kDa Nrf2 form is expressed in all tested cells**. (A) Western blot analysis of the two most abundant Nrf2 forms in lung cancer cell lines H1299, RERF and A549 and in patients’ samples (NSCLC 1 and 2) and normal lung fibroblasts (NLF). Nrf2 was detected with Abcam EP1808Y antibodies. Actin was used as a loading control. (B) Western blot analysis of two most abundant Nrf2 forms in NSCLC cells. Nrf2 was detected with Abcam EP1808Y and Cell signaling (D1Z9C) antibodies. Actin was used as a loading control.

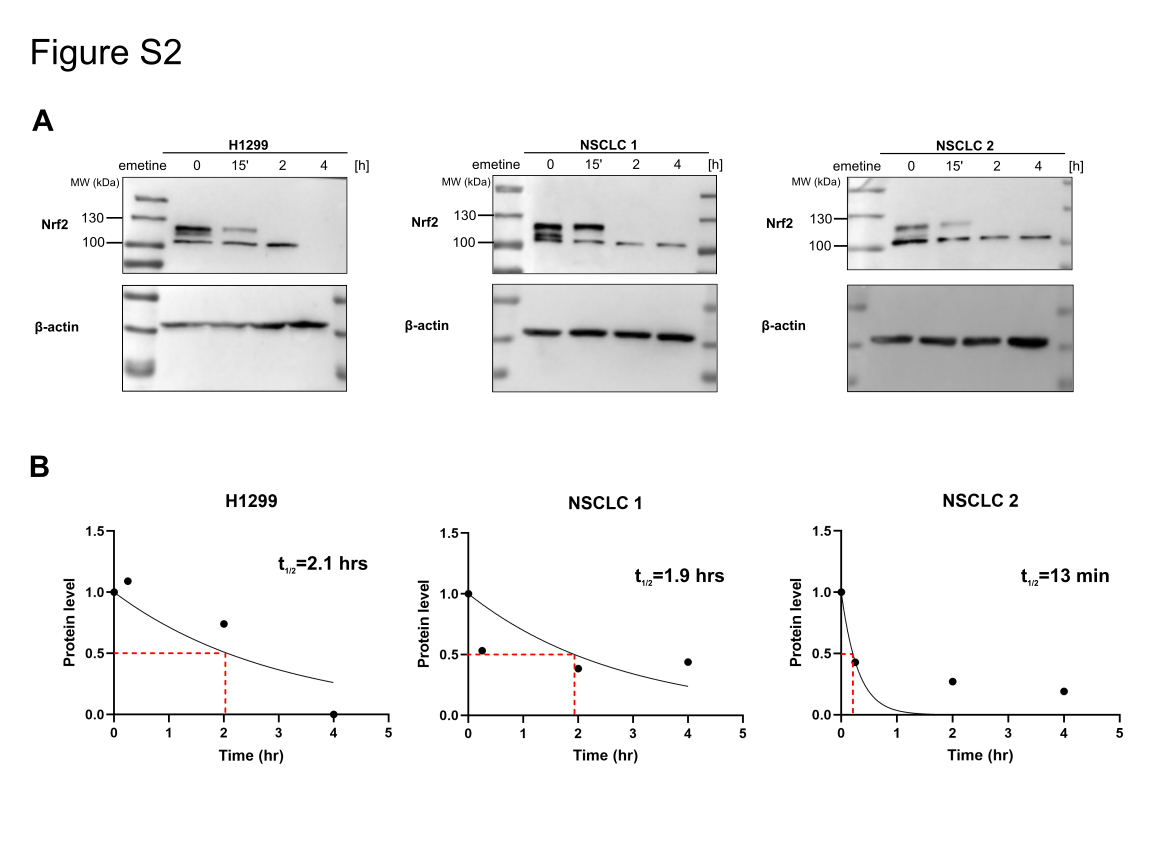
Figure S2. **The stability of newly identified, shorter Nrf2 form after translation inhibition.** (A) Western blot analysis after treatment with translation elongation inhibitor emetine dihydrochloride (20 µM) at indicated time points in H1299, NSCLC 1 and NSCLC 2. Actin was used as a loading control. (B) Curves show data fitted using nonlinear regression and a one-phase exponential decay equation used for calculating the half-life of protein based on the average band densities for each time point, that were normalized to time point zero.

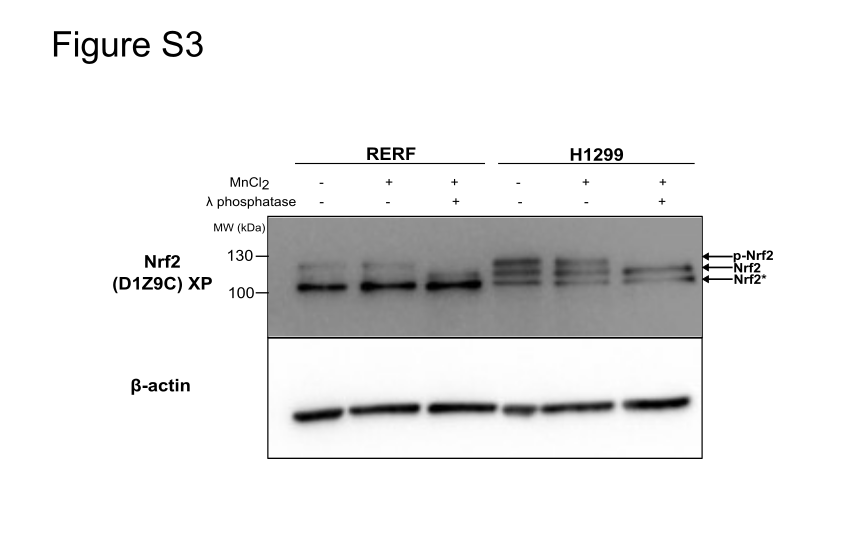
Figure S3. **Stable Nrf2 form is not phosphorylated.** Lambda protein phosphatase (λPP) treatment of RERF and H1299 lysates. Cell lysates were incubated with or without λ phosphatase for 30 min in 30 °C in the presence of MnCl2. Arrows indicate different Nrf2 forms: **←**P for phosphorylated, canonical Nrf2; ← not phosphorylated, canonical Nrf2; **←*** non-canonical Nrf2. Nrf2 was detected by western blot analysis using Cell signaling (D1Z9C) XP antibodies.

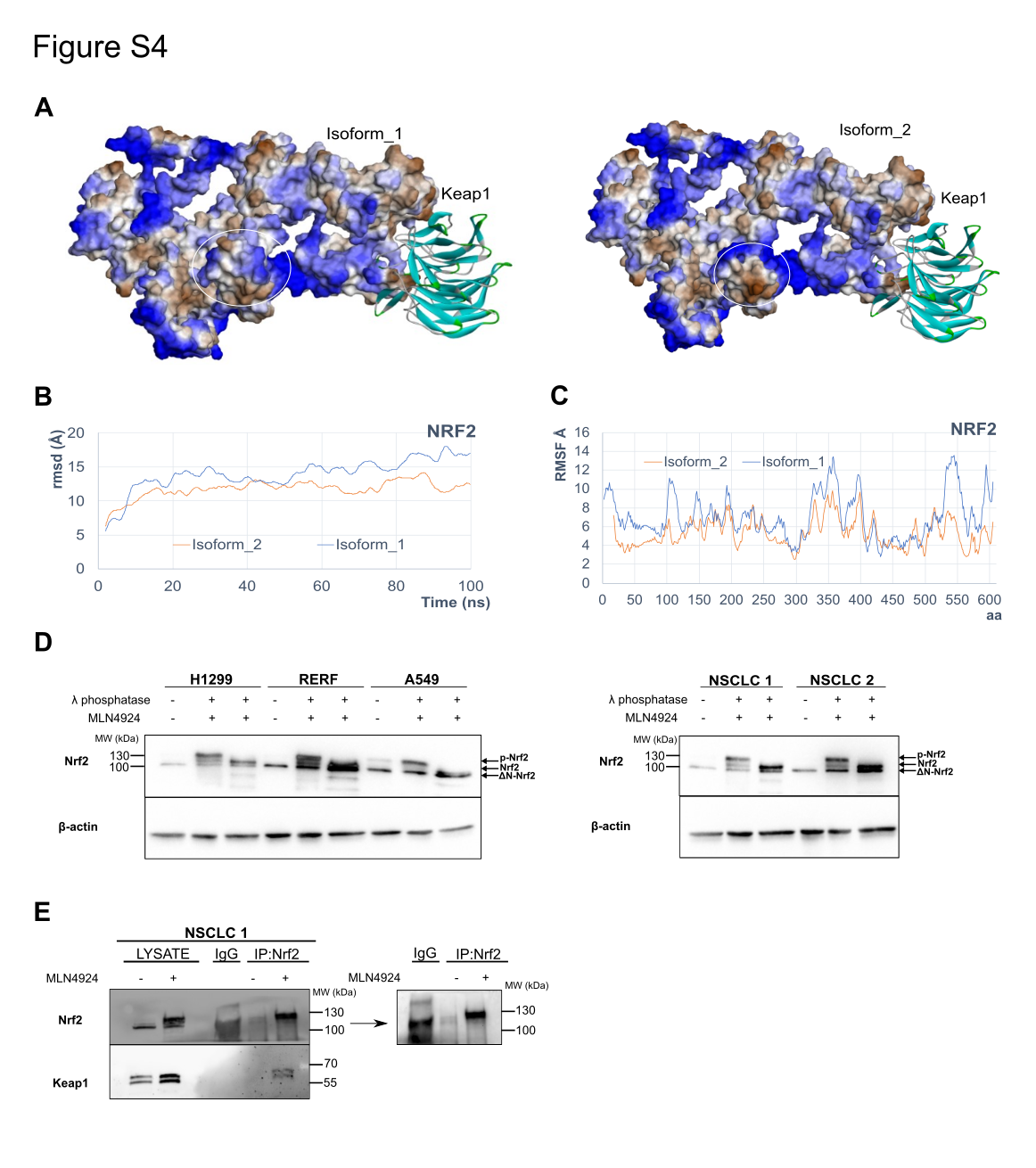
Figure S4. **Binding of ΔN-Nrf2** **to** **Keap1 is impaired.** (A) Left image represents a structure of full-length Nrf2 isoform 1 protein bound to the Keap1, while the image on the right represents a structure of ΔN-Nrf2 isoform 2 protein bound to the Keap1. White circles represent  the difference in the structure of isoform 1 and isoform 2, due to the deletion of the first 16 amino acids. (B) In order to compare the stability of full-length Nrf2 and ΔN-Nrf2, the RMSDs (root-mean-square deviations) were computed. RMSD calculations measure the difference between the backbones of a protein from its initial structural conformation to its final position. The stability of the protein relative to its conformation can be determined by the deviations produced during the course of its simulation (1). A RMSD value is expressed in Ångström (Å) which is equal to 10^−10^ m. (C) RMSF stands for root mean square fluctuation; it calculates the flexibility of individual residues or how much a particular residue fluctuates during a simulation (2). It indicates structurally which amino acids in a full-length Nrf2 isoform 1 and ΔN-Nrf2 isoform 2 protein contribute the most to a molecular motion. (D) Western blot analysis of Nrf2 in lung cancer cell lines H1299, RERF and A549, and in NSCLC cells, after treatment with neddylation inhibitor MLN4924. Cells were treated with 1 μM MLN4924 for 12 h. After treatment with MLN4924, cell lysates were incubated with or without λ phosphatase for 30 min in 30 °C in the presence of MnCl_2_ and Nrf2 was detected by Western blot with Abcam EP1808Y antibodies. Actin was used as a loading control. Arrows indicate different Nrf2 forms: ←p-Nrf2 for phosphorylated, canonical Nrf2; ←Nrf2 for non-phosphorylated, canonical Nrf2; ←ΔN-Nrf2 for N-terminally truncated Nrf2 isoform 2. (E) Western blot analysis of NSCLC 1 cells after co-immunoprecipitation of Nrf2, under homeostatic conditions and after treatment with neddylation inhibitor MLN4924 for 12 hours. Precipitates were probed with anti-Keap1 antibodies to analyze levels of Nrf2-bound Keap1. IgG was used as a control for co-immunoprecipitation. Right side panel presents Western blot results in co-immunoprecipitation of Nrf2 and IgG samples after longer exposure.

**Table S1.** Expression of Nrf2 isoform and loading control (β actin) in H1299 cell line after treatment with translation elongation inhibitor, emetine dihydrochloride (20 µM), at different time points (15 min, 2 and 4 hours).

| **Time points [h]** | **Nrf2 isoform signal** | **Loading control signal** | **Normalization (sample/loading control)** | **Normalization (sample/control sample)** |
| --- | --- | --- | --- | --- |
| 0 | 8321.104 | 10283.25 | 0.8092 | 1 |
| 0.25 | 7008.054 | 7940.004 | 0.8826 | 1.0908 |
| 2 | 9036.004 | 15087.22 | 0.5989 | 0.7401 |
| 4 | 0 | 18733.02 | 0 | 0 |

**Half-life of Nrf2 isoform (best-fit value): 2.067 hours**

**Table S2.** Expression of Nrf2 isoform and loading control (β actin) in NSCLC 1 after treatment with translation elongation inhibitor, emetine dihydrochloride (20 µM), at different time points (15 min, 2 and 4 hours).

| **Time points [h]** | **Nrf2 isoform signal** | **Loading control signal** | **Normalization (sample/loading control)** | **Normalization (sample/control sample)** |
| --- | --- | --- | --- | --- |
| 0 | 9476.347 | 17315.116 | 0.5473 | 1 |
| 0.25 | 6238.69 | 21434.551 | 0.2911 | 0.5318 |
| 2 | 4603.861 | 21916.622 | 0.2101 | 0.3838 |
| 4 | 5730.418 | 23930.986 | 0.2395 | 0.4375 |

**Half-life of Nrf2 isoform (best-fit value): 1.936 hours**

**Table S3.** Expression of Nrf2 isoform and loading control (β actin) in NSCLC 2 after treatment with translation elongation inhibitor, emetine dihydrochloride (20 µM), at different time points (15 min, 2 and 4 hours).

| **Time points [h]** | **Nrf2 isoform signal** | **Loading control signal** | **Normalization (sample/loading control)** | **Normalization (sample/control sample)** |
| --- | --- | --- | --- | --- |
| 0 | 11203.52 | 12061 | 0.9289 | 1 |
| 0.25 | 5413.104 | 13628.49 | 0.3972 | 0.4276 |
| 2 | 3536.276 | 14047.49 | 0.2517 | 0.2710 |
| 4 | 4129.154 | 23391.89 | 0.1765 | 0.1900 |

**Half-life of Nrf2 isoform (best-fit value): 0.2075 hours**

**Table S4.** Sequences of primers for detection of Nrf2 transcript variants in RT-PCR

| **Nrf2 transcript variants** | **Primer sequences** | **Product lengths** |
| --- | --- | --- |
| Transcript 1,6,7 | For: TCATGATGGACTTGGAGCTG  Rev GCAATGAAGACTGGGCTCTC | Tr 1: 475 bp  Tr 6: 385 bp  Tr 7: 256 bp |
| Transcript 6,7 | For: CGACCTTCGCAAACAACTCT  Rev: TGACCGGGAATATCAGGAAC | Tr 6: 817 bp  Tr 7: 688 bp |
| Transcript 2,3,4,5,8 | For: TCCTGCTTTATAGCGTGCAA  Rev: GCAATGAAGACTGGGCTCTC | Tr 2: 602 bp  Tr 3: 581 bp  Tr 4: 476 bp  Tr 5: 568 bp  Tr 8: 531 bp |

**Table S5**. Primer sequences for evaluation of Nrf2 transcript variants expression via RT-qPCR

| **Nrf2 transcript variant** | **Primer sequences** | **Product length** |
| --- | --- | --- |
| 1,6,7 | For:  AACACACGGTCCACAGCTC  Rev: TCTTGCCTCCAAAGTATGTCAA | 102 bp |
| 2,3,8 | For:  GACGGGATATTCTCTTCTGTGC  Rev: CATACTCTTTCCGTCGCTGA | 128 bp |
| 1,2,3,4,5,6,7,8 | For:  GAGAGCCCAGTCTTCATTGC  Rev TGCTCAATGTCCTGTTGCAT | 104 bp |
